# A network-based approach to eQTL interpretation and SNP functional characterization

**DOI:** 10.1101/086587

**Authors:** M. Fagny, J.N. Paulson, M.L. Kuijjer, A.R. Sonawane, C.-Y. Chen, C.M. Lopes-Ramos, K. Glass, J. Quackenbush, J. Platig

## Abstract

Expression quantitative trait locus (eQTL) analysis associates genotype with gene expression, but most eQTL studies only include *cis*-acting variants and generally examine a single tissue. We used data from 13 tissues obtained by the Genotype-Tissue Expression (GTEx) project v6.0 and, in each tissue, identified both *cis*- and *trans*-eQTLs. For each tissue, we represented significant associations between single nucleotide polymorphisms (SNPs) and genes as edges in a bipartite network. These networks are organized into dense, highly modular communities often representing coherent biological processes. Global network hubs are enriched in distal gene regulatory regions such as enhancers, but are devoid of disease-associated SNPs from genome wide association studies. In contrast, local, community-specific network hubs (core SNPs) are preferentially located in regulatory regions such as promoters and enhancers and highly enriched for trait and disease associations. These results provide help explain how many weak-effect SNPs might together influence cellular function and phenotype.

## Introduction

More than a decade after the sequencing of the human genome, our understanding of the relationship between genetic variation and complex traits is limited. Genome Wide Association Studies (GWASes), looking for association between common genetic variants and traits from height [Liu et al., 2010, Wood et al., 2014] to cancer [Chang et al., 2014] to metabolic diseases [Rueedi et al., 2014], have resoundingly shown that complex phenotypes are influenced by many variants of relatively weak effect [Visscher et al., 2012, Stranger et al., 2011]. The failure of GWAS to find common SNPs with large effect sizes for complex phenotypes has led to the speculation that a limited number of rare variants may be driving observed trait variation. However, recent whole genome and exome sequencing in type 2 diabetes, analyzing more than 100,000 individuals, found not only that rare (MAF *<* 0.5%) variants with large effect sizes are unlikely to account for the missing trait heritability in that disease, but the overwhelming majority of variants associated with type 2 diabetes were common [Fuchsberger et al., 2016].

In surveying the single nucleotide polymorphisms (SNPs) in the NHGRI-EBI GWAS catalog, nearly all (*∽* 93%) lie in non-coding regions of the genome [Ward and Kellis, 2012, Tak and Farnham, 2015] and a growing body of work [Maurano et al., 2012, Ernst et al., 2011] has found that GWAS variants are enriched for regions of the genome that affect gene regulation both locally and distally. These results suggest that rather than being driven by a few causative changes in protein coding regions, most complex traits are manifest through complex regulatory effects by many common variants that likely alter gene expression.

Expression Quantitative Trait Locus (eQTL) analysis estimates the influence of a SNP’s presence on the expression of a gene, and most published studies have limited their analysis to *cis*-regulation by focusing only on SNPs in a narrow window around a gene’s transcription start site (TSS). However, there is mounting evidence of the importance of distal or *trans*-regulatory elements as drivers of disease phenotypes [Westra et al., 2013, Kirsten et al., 2015, Franz´en et al., 2016]. As large cohort studies become more frequent [The GTEx Consortium, 2015, Westra and Franke, 2014], allowing for the detection of large numbers of *cis*- and *trans*-eQTLs, new methods will be needed to identify regulatory patterns in the complex web of interactions between common variants and gene expression.

Here we present a systems genetics approach that organizes *cis*- and *trans*-eQTLs as a bipartite network, where each significant SNP-gene association is cast as an edge the network, and shows that the structure of this network provides insight into the collective regulatory role of common variants and the genes they influence. We used genotype and expression data from 13 tissue types obtained by the Genotype-Tissue Expression (GTEx) project v6.0 [The GTEx Consortium, 2015] and, in each tissue, identified both *cis*- and *trans*-eQTLs to create a tissue-specific network. We find that these networks possess substantial community structure, meaning that SNP and gene nodes are organized into densely connected groups or communities. These communities often comprise groups of functionally related genes from multiple chromosomes, with these genes organized around the SNPs that influence their expression.

The functionally related genes in these communities are often tissue-specific and related to processes important for that tissue. For example, we find communities enriched for contraction and ventricular cardiac muscle tissue morphogenesis in heart tissue, and nerve impulse and myelination in tibial nerve tissue, among others. Those SNPs which are central to their communities are not only more likely to lie in active regions of the genome, but are also significantly enriched for GWAS associations. These results are highly consistent across tissue types, suggesting that a network representation of *cis-* and *trans-*eQTLs provides a natural framework for understanding the collective impact of genetic variants on gene regulation and disease pathogenesis.

## Results

### eQTL mapping

We downloaded the Genotype-Tissue Expression (GTEx) version 6.0 data set (phs000424.v6.p1, 2015-10-05 release) from dbGaP (approved protocol #9112). For the RNA-Seq data, we used the Bioconductor R YARN package [Paulson et al., 2016b] to perform quality control, gene filtering, and normalization preprocessing [Paulson et al., 2016a]. This process included the identification and removal of GTEX-11ILO due to potential sex misannotation, as well as filtering and normalization of the data in a tissue-aware manner using smooth quantile normalization [Hicks et al., 2016] and removed sex-chromosome and mitochondrial genes (retaining 29,242 genes; see Methods for details of tissue-specific gene filtering). We also grouped skin samples from the lower leg (sun exposed) and from the suprapubic region (sun unexposed) based on gene expression similarity. For our analysis we only considered tissues for which we had both RNA-seq and imputed genotyping data for at least 200 individuals. After all preprocessing steps, 13 tissues met all criteria (Figure 1 and Table 1).

**Figure 1:**
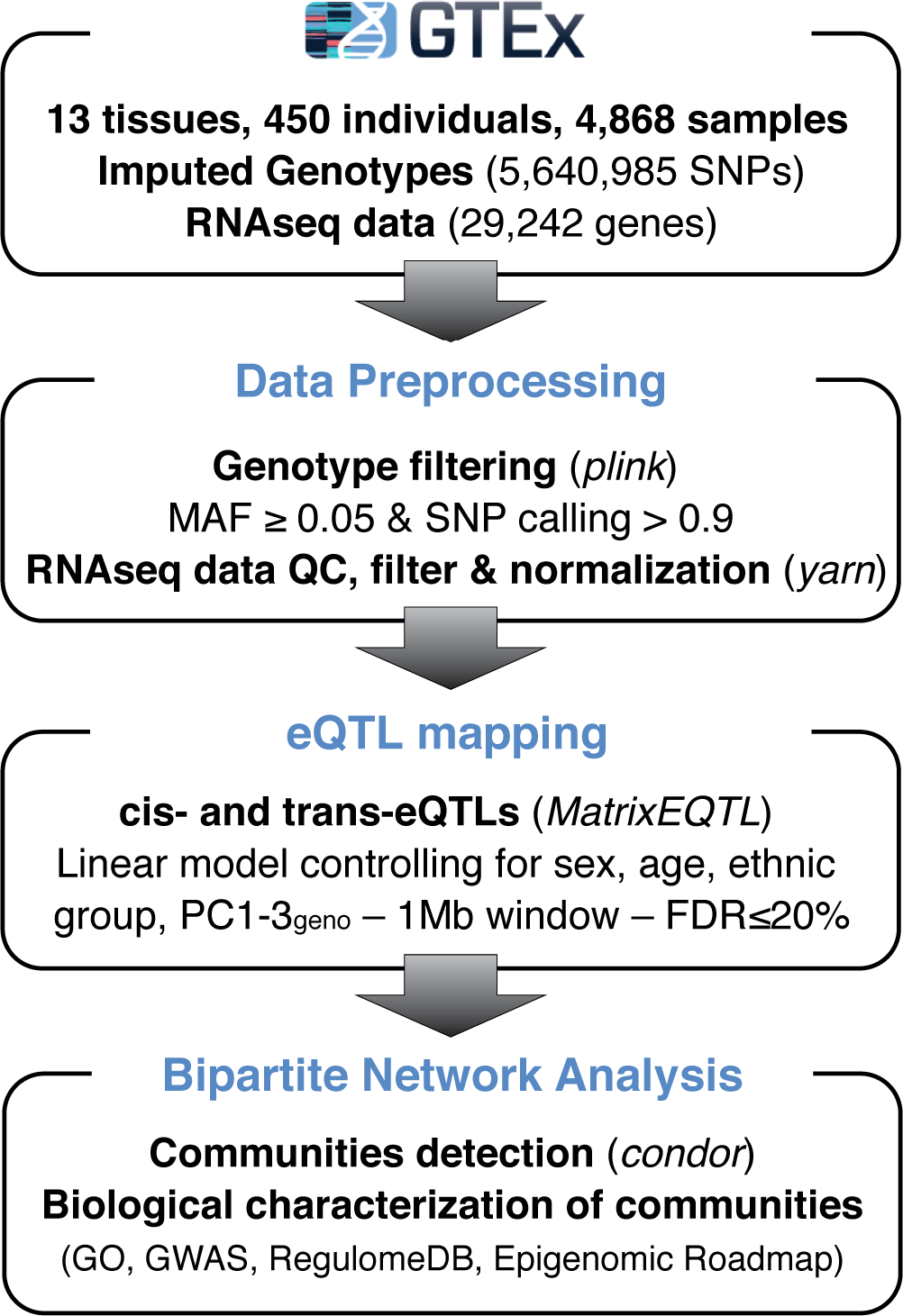
Study overview. Principal components 1-3 used as covariates in eQTL mapping are presented in Supplementary Figure S1. Comparison of our eQTL mapping results with those from the GTEx consortium are presented in Supplementary Table S1

**Table 1:** Summary of data: tissues and number of samples.

| **Tissue names** | **Abbreviation** | **GTEx tissues** | **Samples** |
| --- | --- | --- | --- |
| Adipose subcutaneous | ADS | Adipose - Subcutaneous | 313 |
| Aorta | ATA | Artery - Aorta | 216 |
| Artery tibial | ATT | Artery - Tibial | 303 |
| Fibroblast | FIB | Skin cells - Transformed fibroblasts | 291 |
| Esophagus mucosa | EMC | Esophagus - Mucosa | 276 |
| Esophagus muscularis | EMS | Esophagus - Muscularis | 245 |
| Heart left ventricle | HRV | Heart - Left ventricle | 212 |
| Lung | LNG | Lung | 290 |
| Skeletal muscle | SMU | Muscle - Skeletal | 378 |
| Tibial nerve | TNV | Nerve - Tibial | 278 |
| Skin | SKN | Skin - Not sun exposed (Suprapubic) | 127 |
|  |  | Skin - Sun exposed (Lower leg) | 243 |
|  |  | Total | 370 |
| Thyroid | THY | Thyroid | 295 |
| Whole blood | WBL | Whole blood | 365 |

For each of the 13 tissues, we tested for association between SNP genotypes and gene expression levels both in *cis* and in *trans*, correcting for reported sex, age, ethnic background and the top 3 principal components obtained using genotyping data (Supplementary Figure S1, Figure 1, and Methods). Including SNPs within a +/−1Mb window around each gene, we found between 285,283 and 691,333 significant *cis*eQTLs (5,301 – 11,035 genes) and between 7,151 and 15,183 significant *trans*-eQTLs (326 – 955 genes) at a FDR of 5% in each tissue. Consistent with previous studies [Dimas et al., 2009, Stranger and De Jager, 2012, Veyrieras et al., 2008], we find most *cis*-eQTLs are located around TSSes, with 50% of SNPs located within *~* 16,000bp of the nearest TSS (14,767 bp for whole blood – 17,109 bp for thyroid). We also observed that *cis*-eQTLs were highly replicable across tissues (70–88% were replicated in at least one other tissue), while *trans*-eQTLs replicability across tissues was more varied (37–88% were replicated in at least one other tissue). A majority of *trans*-eQTLs were also associated with genes in *cis* (50% – 66%).

To compare our results with those reported by the GTEx consortium, we downloaded the single tissue *cis*-eQTLs from the GTEx portal (http://www.gtexportal.org/). Despite differences in normalization, number and type of covariates, and eQTL p-value calculation, at a FDR of 5%, on average 73% of our cis-eQTLs were also detected by the GTEx project in each tissue (minimum 70% in artery aorta and maximum 76% in thyroid, Supplementary Table S1).

### eQTL networks are highly modular

For each of the 13 tissues, we represented the significant eQTLs as a bipartite network, with nodes representing either SNPs or genes, and edges representing significant SNP-gene associations (see, for example, heart left ventricle tissue Figure 2A). Each network was comprised of a giant connected component (GCC) plus additional small connected components. To increase the size of the GCC and because network centrality measures are more sensitive to false negative than to false positive edges [Platig et al., 2013, Wang et al., 2012], we relaxed the FDR cut-off and included all eQTLs with FDR q-values under 0.2. The resulting 13 tissue-specific eQTL networks had GCCs which included 160,634 – 812,141 edges, 50,999 – 401,149 SNPs, and 1,138 – 9,492 genes (Figure 2B-D); we focused on the networks defined by these GCCs for all subsequent analysis.

**Figure 2:**
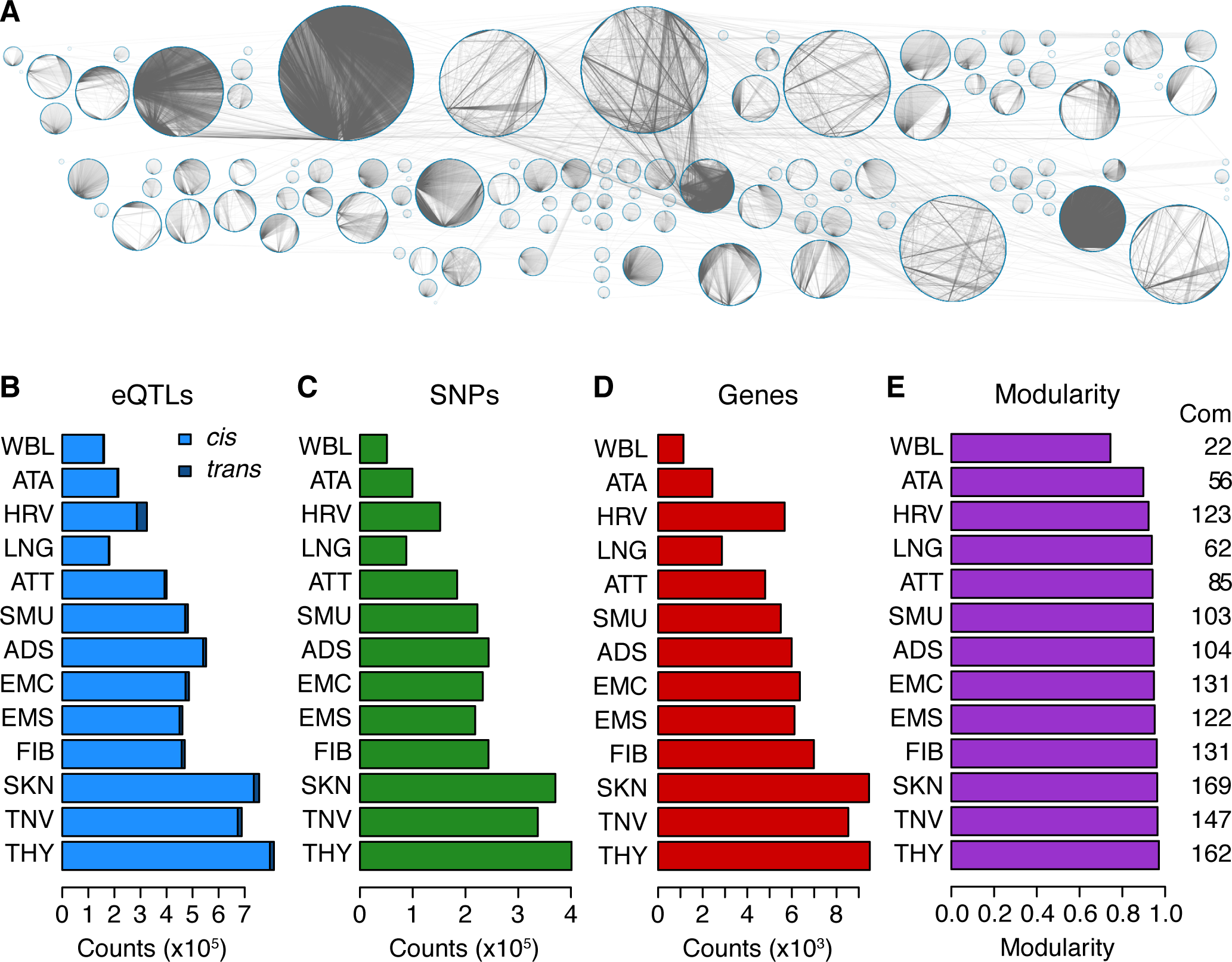
Structure of eQTL networks. **A.** The eQTL network from heart left ventricle, each circle represents a community. **B.** Number of eQTLs in each network. **C.** Number of SNPs in each network. **D.** Number of genes in each network. For a comparison between genes in eQTL communities and clusters derived from hierarchical clustering of gene expression, see Supplementary Figure S4. **E.** Modularity of the eQTL network from each of the thirteen tissues. The number of communities (Com) in each network is written on the right side of the figure.

We previously reported that bipartite eQTL networks are organized into highly modular communities, each of which is comprised of SNPs and genes with a greater number of significant associations than expected given the distribution of node degrees [Platig et al., 2016]. Motivated by our previous findings, we used the R condor package [Platig et al., 2016] to identify communities in each of the 13 eQTL networks (Figure 1). We found between 22 (whole blood) and 169 (skin) communities in each tissue-specific network. The modularities of the eQTL networks ranged from 0.74 (whole blood) to 0.97 (thyroid, Figure 2E). This means that within each network, the communities were well defined, with the density of edges linking nodes from the same community being much higher than the density of edges linking nodes from different communities (see example in Figure 3A, Supplementary Figure S3). The size of the communities ranged broadly, between 2 and 1,220 genes and between 3 and 26,056 SNPs.

**Figure 3:**
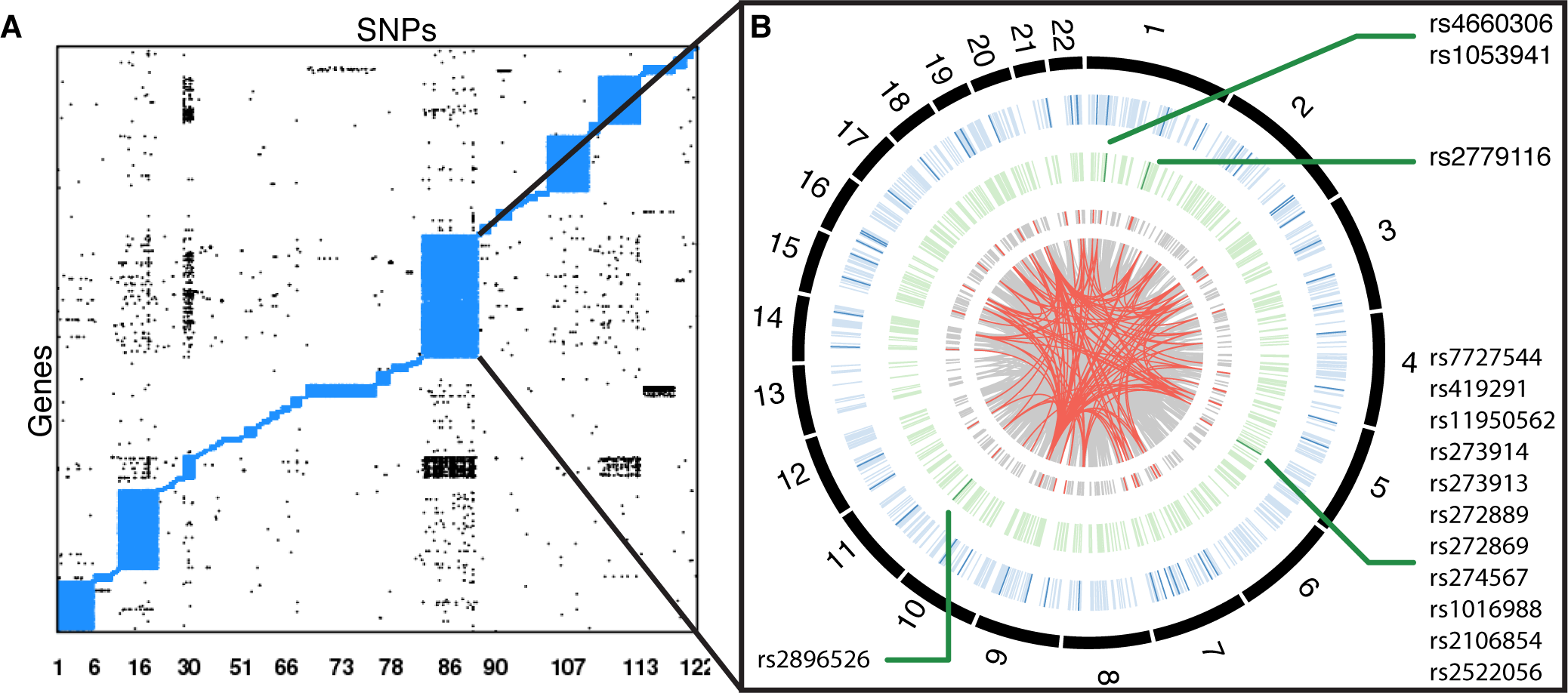
Close-up of the heart ventricle community 86. **A.** Structure of communities within the eQTL heart left ventricle network. Each network edge is represented by a point. Intra-communities edges are plotted in blue, inter-community edges in black. Community structure for the other twelve networks are presented in Supplementary Figure S3. **B.** Circos diagram for heart left ventricle eQTLs in community 86. All SNP-gene assocations from the eQTL analysis are shown as grey lines, with *trans*-eQTLs around innermost circle and *cis*-eQTLs around the second circle. Red lines represent eQTLs linked to genes involved in cellular respiration or SNPs involved in metabolism. SNPs are shown as green bars in the third circle, with dark green bars indicating an association with metabolic traits. Details about GWAS annotation of the SNPs are given in Supplementary Table S3. The fourth circle (blue bars) displays genes from each eQTL association. Dark blue bars represent genes associated to cellular metabolism. Details about GWAS annotation of the SNPs and genes are given in Supplementary Table S3. A barplot summarizing the number of chromosomes present in each community of each network is shown in Supplementary Figure S2.

We recognize that genetic recombination effects, including local linkage disequilibrium, might lead to a clustering of SNPs and genes from the same chromosome (or chromosomal region) to cluster together. To test this, we examined the chromosomal distribution of genes and SNPs in each community. Across all 13 tissues, between 71% (fibroblast cell line) and 94% (adipose subcutaneous) of communities included genes and SNPs from two or more chromosomes (Supplementary Figure S2). For example, Figure 3 shows significant associations between SNPs and genes from all 22 autosomes in community 86 of heart left ventricle.

To determine how much correlated gene expression determined community structure, we performed an average linkage hierarchical clustering using a Pearson correlation metric for the expression levels of genes in each network. For each tissue, we set the distance threshold to select as many groups of genes in the clustering dendrogram as there were communities. We used the Jaccard Index to compare the genes in hierarchical clustering groups with those in the network communities. While some small communities in each network have highly correlated expression (maximum Jaccard Index of 0.81 for one community in heart left ventricle), gene expression correlation is not the main driver of the eQTL communities (Jaccard Index median of 0.04 and 95^th^ percentile of 0.14 across all tissues, Supplementary Figure S4). This suggests that there may be other, biologically relevant factors driving the structure of the eQTL network communities.

### Communities cluster genes by biological functions

We tested the communities in each tissue for over-representation of Gene Ontology (GO) biological processes [Ashburner et al., 2000]. Across all tissues, we found 208 communities enriched for at least one biological process at a FDR of 5% (Supplementary Table S2). We compared the occurrence of biological processes across communities and tissues, and found that some communities were enriched for genes involved in tissue-specific functions while others included genes with biological functions relevant to all tissues (Figure 4).

**Figure 4:**
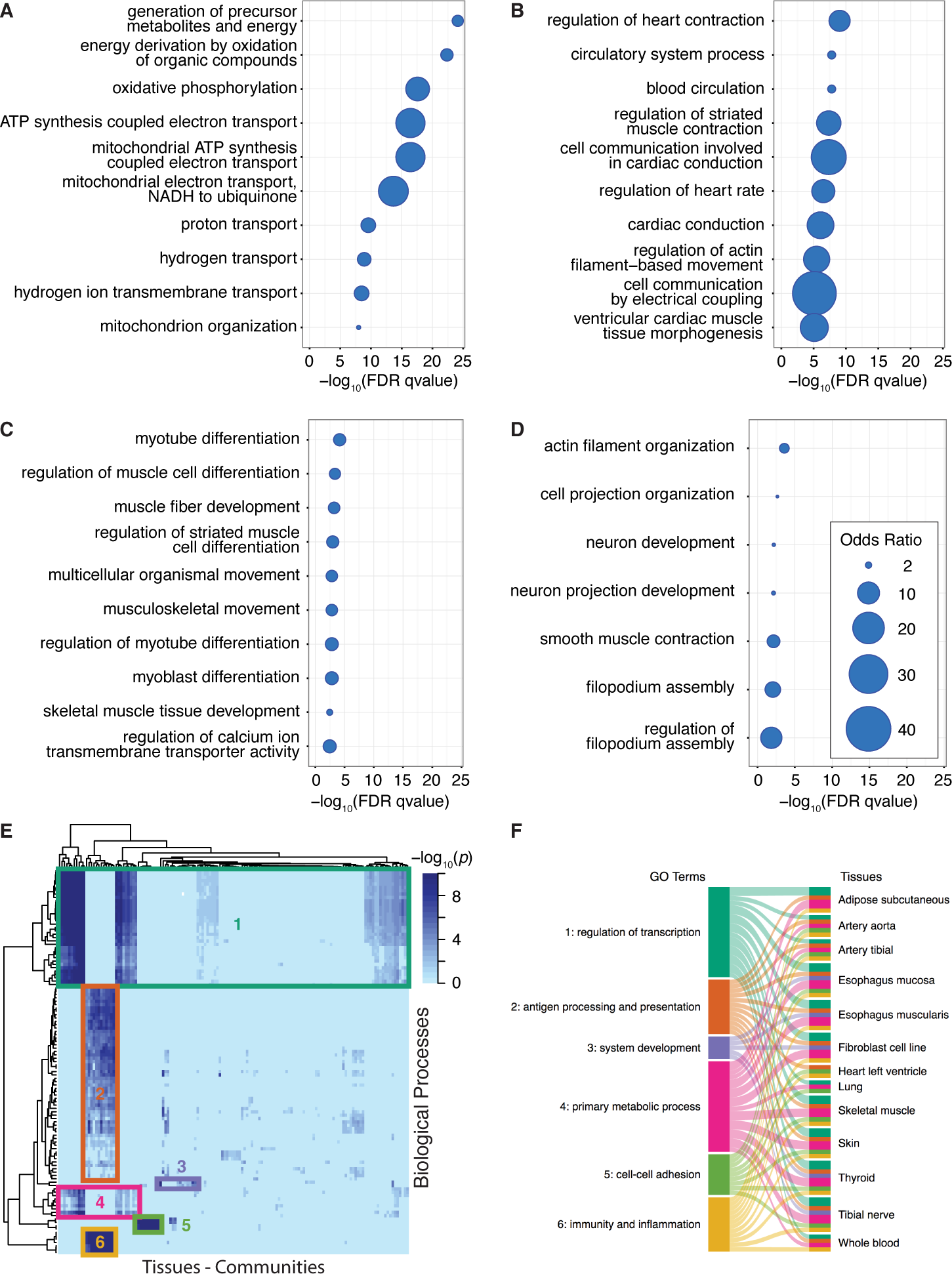
Network communities are enriched for specific biological functions. A complete list of the significantly over-represented biologic processes in each community and each network can be found in Supplementary Table S2. **A.** Heart left ventricle community 86 is enriched for genes involved in cellular respiration. **B.** Heart left ventricle community 30 is enriched for genes involved in cardiac muscle development and contraction. **C.** Muscle skeletal community 75 is enriched for genes involved in striated muscle development. **D.** Esophagus muscularis community 83 is enriched for genes involved in smooth muscle contraction. **E.** Heatmap clustering similarity of GO biological processes in communities from all tissues. Only GO terms that were significant in at least ten tissues were included. **F.** Sankey diagram linking each clusters from the heatmap to the tissues that contain at least one community enriched for genes involved in these functions. A ribbon’s thickness is proportional to the number of communities enriched for each cluster of GO terms in each tissue-specific network.

We used the GO enrichment analysis to identify tissue-specific functions, which we defined as a community-enriched GO term appearing in only a single tissue. An interesting example is community 86 from the heart left ventricle network (Figure 4A). This community is enriched for genes involved in cellular respiration and the mitochondrial respiratory chain (Supplementary Table S2). These functions, despite being ubiquitous, are particularly important in heart due to its high metabolic requirements, and studies have shown that dysfunction of the respiratory chain is involved in many heart diseases [Schwarz et al., 2014]. This heart community contains 13,143 SNPs linked to 1,182 genes (grey links, Figure 3B). Of these, there is a subset involved in cellular respiration (red links) that includes 560 SNPs linked to 52 genes, located on 20 autosomes. GWASes have reported 9 of these 52 genes to be associated with cardiovascular-related traits and metabolism such as conotruncal heart defects, obesity and blood metabolite levels (Figure 3B, Supplementary Table S3, [Welter et al., 2014]). This community also contains SNPs from three genomic regions that have been linked to metabolic traits and blood metabolite levels; this includes association with the metabolite carnitine, a molecule involved in the transport of fatty acid from cytoplasm to the mitochondrial matrix where those fatty acids are metabolized (Figure 3B, Supplementary Table S3).

Other examples of tissue-specific over-representation of biological functions include transmission of nerve impulse and myelination in community 4 from tibial nerve, and muscle development and contraction in muscular tissues. Community 30 from the heart left ventricle network is enriched for genes related to ventricular cardiac muscle tissue morphogenesis and contraction (Figure 4B), community 75 from skeletal muscle for striated muscle contraction and cell differentiation (Figure 4C), and community 83 from esophagus muscularis for smooth muscle contraction (Figure 4D and Supplementary Table S2).

We also identified communities enriched for ubiquitous biological functions, which we defined as GO terms which were enriched in a community from at least 10 (of the 13) tissue-specific eQTL networks. To analyze these results, we clustered communities using their enrichment p-values in these ubiquitous biological functions only (Figure 4E). We identified 6 groups of GO terms related to regulation of transcription and RNA metabolism, immunity, system development and cell-cell adhesion that defined 6 groups of communities (Figure 4E-F). For example, we found that genes involved in functions linked to pathogen recognition, innate and adaptive immune response triggering and inflammation (groups 2 and 6, Figure 4E-F and Supplementary Table S2) are over-represented in at least one community in all tissue-specific eQTL networks except lung. These communities include many genes from the major histocompatibility complex (*MHC*) class II and may reflect the presence of infiltrated macrophages in all tissues. Communities within each group in the heatmap share many of the same eQTLs despite their different tissue of origin and the inclusion of their own unique, tissue-specific SNPs and genes (average pairwise Jaccard index of 0.46 between communities groups).

Another category of genes, related to the control of transcription and RNA metabolism, is strongly enriched in communities appearing in 12 of the 13 tissue-specific networks (groups 1 and 4, Figure 4E-F and Supplementary Table S2). In contrast to communities enriched in immune response, these communities do not overlap in terms of eQTLs (average pairwise Jaccard index of 0.03).

This supports our previous finding that eQTL network community structure reflects many SNPs working together to influence groups of functionally related genes and that the communities capture unique features of tissues and phenotypic states. It should be noted, however, that although we find communities in many tissues that have the same “functional enrichment,” this does not necessarily imply that those communities contain identical sets of SNPs, genes, or eQTLs. Rather, although there are varying degrees of overlap depending on the process involved, each tissue has its own combination of SNPs and genes that contribute to supporting processes shared with other tissues. These differences may be due to the activity of distinct biological pathways in each tissue, but we cannot exclude the possibility that they are due to the small differences in the populations analyzed for each tissue. Regardless of the cause, our analysis of these eQTL communities paints a robust picture of communities that involve functions shared across tissues as well as those that are highly specialized to individual tissues.

### Network centrality reflects SNP regulatory roles

A number of studies have shown that eQTL SNPs are over-represented in regulatory regions [Battle et al., 2015, The GTEx Consortium, 2015, Ward and Kellis, 2012], but not all eQTL SNPs have obvious regulatory roles. One natural hypothesis that emerges from our structural analysis of eQTL networks is that there may be network properties that are associated with the regulatory roles of SNPs. Specifically, one might expect that SNPs that are more highly connected, either globally or locally, would have different regulatory roles than SNPs at the periphery of the network.

We investigated the relationship between eQTL network structure and regulatory role using two measures of centrality: the degree—the total number of genes to which each SNP is linked—and the modularity contributed by each SNP or “core score” (see Eq. 2 in Methods). A SNP’s degree is simply number of edges associated with that SNP and reflects the global SNP centrality within the entire network, while the core score reflects the local centrality of each SNP within its community.

For each tissue-specific network, we studied the enrichment of central SNPs in eight active regulomeDB SNP categories (A-H vs. I, cf. definitions in Table S4), using all SNPs in the giant connected component as background and correcting for gene density (see Methods). In 10 of the 13 tissues we found that both global hubs (SNPs with degree greater than or equal to 10) and core SNPs (SNPs with a core score in the 4^th^ quartile) were significantly enriched in open chromatin regions, with higher than expected representation for DNase I or transcription factor (TF) binding sites (categories F and G in Figure 5A,B and Supplementary Table S5). Moreover, in 11 of the 13 tissues, core SNPs showed enrichment in regulatory regions that are marked by the association of DNase I peak footprint and peaks, TF binding sites and TF motifs (categories A and B, Figure 5A and Supplementary Table S5).

**Figure 5:**
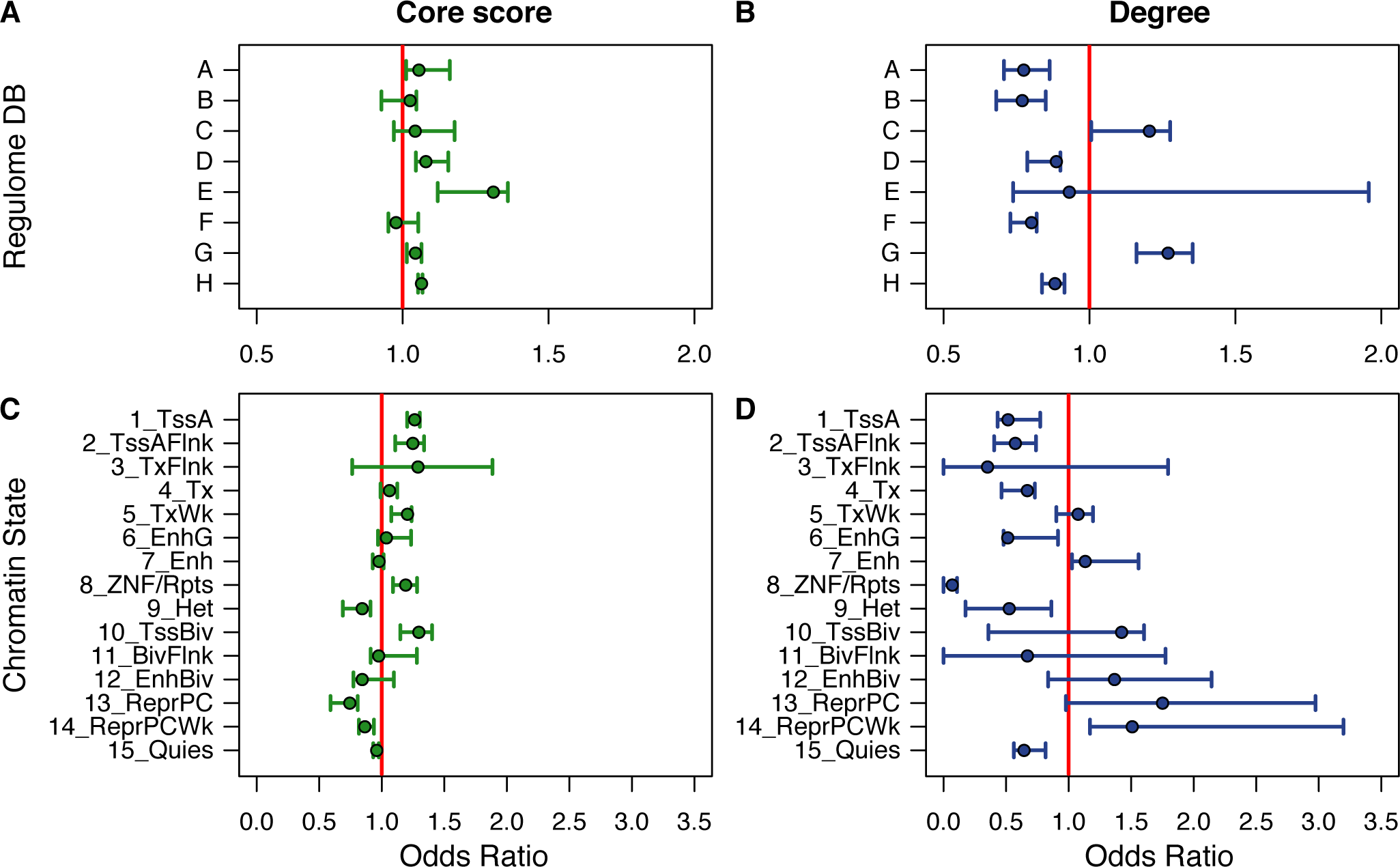
Central SNPs lie in active chromatin regions. Each line shows the distribution of odds ratios (medians as circles, interquartile ranges as bars) for all 13 tissues. Each odds ratio measures the enrichment of central SNPs in a particular functional category, corrected for number of genes within 1Mb of the SNP. In rows: (**A, B**) RegulomeDB: enrichment in genomic regions presenting active marks (categories A-H) compared to no active marks (category I). The definitions of regulomeDB categories used here can be found in Supplementary Table S4. Odds ratios and p-values for each tissue-specific network are listed in Supplementary Table S5. (**C, D**) Roadmap Epigenome Project: enrichment in each chromatin state. Odds ratios and p-values for each tissue-specific network are listed in Supplementary Table S6. Enrichment in each chromatin state for all trans-eQTLs are presented in Supplementary Table S7. In columns: (**A, C**): Core-scores: enrichment in each genomic region for SNPs with a core-score in the top 25%. (**B, D**): Degree: enrichment in each genomic region for SNPs with network degree greater than 10.

Because RegulomeDB integrates data from multiple cell types and tissues, some of these results may not capture tissue-specific effects. To test for tissue-specific associations between regulatory state and centrality in each eQTL network, we used the Roadmap Epigenome Project data, which classified the genome into 15 chromatin states based on epigenetic marks measured in a specific tissue or cell line [Roadmap Epigenomics Consortium et al., 2015]. We calculated enrichment of central SNPs (see Methods) in the 8 tissues (adipose subcutaneous, artery aorta, fibroblast cell line, esophagus mucosa, heart left ventricle, lung, skeletal muscle, and whole blood) for which chromatin state maps were available. We found that the global hubs are generally more likely to lie within non-genic enhancers and Polycomb repressed regions (states 7 and 13-14, Figure 5D), while core SNPs are enriched for promoters, transcribed regions, genic enhancers, and ZNF/Repeats (states 1-5, 6 and 8, Figure 5C). As expected, both global and core SNPs are depleted in constitutive heterochromatin and quiescent regions (states 9 and 15).

Overall, we find that global hubs and community cores each have their own distinct association with active regulatory regions: global hubs are preferentially located in distal regulatory regions while core SNPs are overrepresented in proximal regulatory elements. This is another case where the eQTL network structure provides insight into biological processes; an insight that is reproducible across tissues.

### SNPs in community cores are associated with trait and disease phenotypes

We tested whether the SNPs with high centrality were more likely to be associated with complex traits in GWAS by mapping trait-associated SNPs in the NHGRI-EBI GWAS catalog [Welter et al., 2014] to each tissue-specific eQTL network, considering only SNPs with GWAS *P*-values less than 10^−8^ (GWAS SNPs). In each tissue, we compared the distribution of degrees (number of edges per node) for all SNPs and GWAS SNPs. We found that SNPs of low degree (1-2) were significantly depleted in GWAS SNPs while SNPs of intermediate degree (5-10) showed consistent significant enrichment for GWAS SNPs across all 13 tissues and SNPs of degree greater than 15 were devoid of GWAS associations (Supplementary Figure S5).

These results are even more striking when we consider only GWAS SNPs associated with phenotypes related to the tissue of interest. For example, when we mapped SNPs associated with autoimmune diseases to the whole blood eQTL network, we found that they were enriched for intermediate degree (Figure 6A). In our 2016 analysis of COPD eQTL networks in lung tissue, the over-representation of intermediate degree SNPs led us to discover that GWAS SNPs tend to be highly central not on a global-scale, but instead within the eQTL communities. We tested the generalizability of this observation by comparing the core scores for GWAS SNPs from the EBI-NHGRI GWAS catalog to non-GWAS SNPs. Using a likelihood ratio test and controlling for linkage disequilibrium, we found that GWAS and non-GWAS SNPs had significantly different core score distributions, with the median core scores for GWAS SNPs higher in all 13 tissues (Supplementary Figure S6). This enrichment for high core scores was also present when considering only SNPs associated to autoimmune diseases (shown for whole-blood in Figure 6B, *P* = 1.3 *×* 10^−13^).

**Figure 6:**
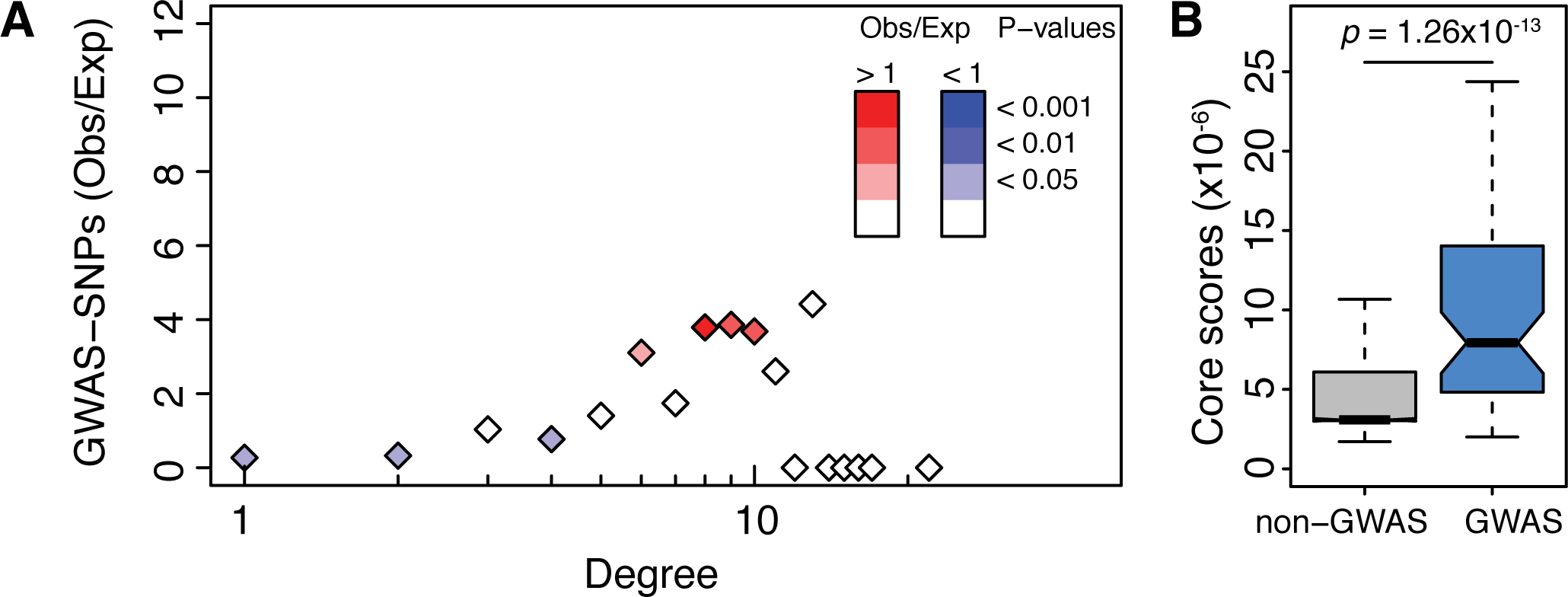
Network properties of GWAS SNPs associated with autoimmune diseases in whole blood. **A.** Ratio of observed vs. expected number of SNPs associated to autoimmune diseases by GWAS studies depending on their network degree. P-values were obtained using 1,000 resamplings taking into account gene density around each SNP. Ratios including all GWAS traits and diseases in each tissue-specific network are in Supplementary Figure S5. **B.** Distribution of core scores for SNPs associated (in blue) or not (in grey) to autoimmune diseases by GWAS. P-values were obtained using a likelihood ratio test and pruning for SNPs in linkage disequilibrium. Distributions for all tissue-specific networks including all GWAS traits and diseases are in Supplementary Figure S6.

These results in the GTEx tissues further support our previous finding that GWAS SNPs have a unique pattern of connectivity within eQTL networks. In each tissue, we find that GWAS SNPs cluster within a relatively small number of communities and within those communities, they are over-represented in local, not global, hubs within the network. These findings are wholly consistent with what one should expect from genotype associations in complex disease. Disease-linked SNPs map to a relatively small number of relevant processes in each tissue, and the likelihood of being disease associated is related to the likelihood that a variant is at the center of its functional community—and therefore more likely to perturb function.

## Discussion

A fundamental tenet of biology is that genotype encodes phenotype. However, the vast majority of traits, including those associated with many diseases, are complex, multifactorial phenotypes in which there is no single, highly penetrant genetic variant. Rather, a growing number of population genetic studies, including GWASes, have shown that complex traits are associated with many (and generally very many) genetic variants of small effect that collectively influence the likelihood that a particular trait or disease will manifest itself.

Expression quantitative trait locus studies associate the genotype at a locus with the expression of genes across the genome. While most eQTL studies have limited their analyses to *cis*-eQTLs, a growing body of literature suggests that *trans*-eQTLs are important in understanding genetic control of expression and that *trans*-eQTLs can help explain disease and other phenotypes. When we examined the presence of *trans*eQTL SNPs among the 15 chromatin states within the 8 tissues annotated by the Roadmap Epigenome Project, we found that *trans*-eQTL SNPs are preferentially located in active regions such as promoters and gene enhancers, while they are significantly depleted in constitutive heterochromatin, repeat, and quiescent regions (Supplementary Table S7). These results support a regulatory role for *trans*-eQTL SNPs and argues for their inclusion in eQTL analyses. The question is then how to understand their collective impact.

Here we present an integrated representation of both *cis*- and *trans*-eQTLs as elements in a bipartite network where a SNP and gene are connected if the genetic variant (SNP) is significantly associated with the gene’s expression. By capturing significant associations between SNPs and the expression levels of genes as elements of a bipartite network, we are able to use the structure of that network and gain insight into complex genetic regulation of gene expression.

Despite the fact that we see significant tissue-specific patterns of gene expression, the global structure of the bipartite eQTL network is highly consistent across all 13 tissues. As we previously reported in COPD[Platig et al., 2016], GWAS SNPs are underrepresented among low degree nodes in the network, overrepresented at intermediate degree, and absent at high degree. This non-random association of GWAS association with eQTL network degree suggests that there is some nontrivial structure in the network that might be associated with the GWAS SNP distribution.

We used a community detection algorithm [Barber, 2007] to search for interrelated communities of SNPs and genes and found that the eQTL network in each of the 13 tissues was organized into a highly modular structure. These eQTL communities suggest that groups of SNPs influence groups of genes. When we look at the genes represented in each community, we find that many communities are enriched for genes with similar functions or are associated with coherent biological processes and span multiple chromosomes. When comparing communities between tissues, we find significant differences in community functional repertoire, consistent with the tissue-specific expression that we observe. We do, however, find many communities with common patterns of functional enrichment shared between tissues where those functions are relevant to the tissues in question.

The eQTL communities have an internal structure with differences in the within-community connectivity of the SNPs. Using a core score to quantify those differences, we find that core SNPs are more likely to be in regulatory regions, more likely to be in active chromatin, and more likely to be associated with traits in GWASes. This is true for both all GWAS SNPs and those with tissue-specific associations. When comparing core SNPs to global network hub SNPs, we find core SNPs are enriched for GWAS associations and active chromatin. We also find that global hubs are associated with distal regulatory gene regions while community core SNPs are concentrated in proximal regulatory regions.

These results paint a compelling picture of how genetic variants can collectively influence traits, including many diseases. We know that regulation of gene expression is a complex process, involving many factors, and the modular structure of the eQTL networks predicts that groups of genetic variants influence collections of genes in the network. Our interpretation of this association with complex traits is that core SNPs are those in the network most likely to produce coherent changes in functionally related genes or biological processes.

Our results also indicate representing eQTLs as a bipartite graph encompassing both *cis*- and *trans*acting SNPs, can help uncover many features of biological systems that are otherwise difficult to discover. We know that there are many nonlinear aspects of the gene regulatory process, such as epistasis, and it may well be that the eQTL bipartite graphs provide a filter that can help us discern the complexity of these interactions.

GWASes have been successful in identifying genes and, in some cases, pathways that are implicated in disease processes [Hirschhorn, 2009]. However, studies with ever-larger populations are not finding variants of large phenotypic effect, but instead identify more and more variants of weak effect [Wood et al., 2014]. Further, there are now more than 16,500 SNPs that have been associated with one or more phenotypic traits [Hall et al., 2016]. With nearly all of these SNPs having small effect sizes, it has been di?cult to interpret the functional link between genotype and phenotype.

When we mapped 97 SNPs associated with autoimmune diseases to the whole blood eQTL network, we found that most (88) fall within 3 communities, with a minority (9) appearing in 8 other communities. The three main communities are enriched for Gene Ontology biological processes that include immune response, inflammatory response and response to stress (Supplementary Table S2). When we examined the genes directly liked to these 97 SNPs via eQTL association, we found the genes were enriched for cell killing, response to bacterium, T-cell co-stimulation, and innate and adaptive immune responses. In the heart ventricle, we found GWAS SNPs associated with metabolism mapped to a community enriched for cellular respiration genes, providing a biologically plausible picture of the way in which these GWAS SNPs could lead to observed phenotypes. This suggests that these eQTL communities group together functionally related sets of variants, including GWAS SNPs, that they do this in a tissue-specific manner, and that the associations can help explain the biology of the tissues and the diseases that affect them.

The work presented here represents a new way of interpreting the link between genetic variation and gene expression. The consistency of the results across 13 tissues suggests that these eQTL networks capture universal features of the way in which genotype influences phenotype, and one that embraces the polygenic nature of complex traits, including disease.

## Methods

### GTEx data preprocessing, filtering, and merging

We downloaded NHGRI GTEx version 6 imputed genotyping data and RNA-seq data from the dbGaP database. The RNA-Seq data were preprocessed using Bioconductor R YARN package [Paulson et al., 2016a, Paulson et al., 2016b] and normalized using Bioconductor R qsmooth package [Hicks et al., 2016]. We excluded five sex-specific tissues (prostate, testis, uterus, vagina, and ovary), and merged the skin samples from the suprapubic and lower leg sites. We limited our eQTL analysis to the 13 tissues for which there were available data for at least 200 individuals. The RNA-seq and genotyping data were mapped by the GTEx Consortium to GENCODE version 19, which was based on human genome build GRCh37.p13 (Dec 2013). We accounted for RNA extraction kit effects using the *removeBatchEffect* function in the R R limma package package [Ritchie et al., 2015].

### eQTL mapping and bipartite network construction

For eQTL analysis, we excluded SNPs from all analyses if they had a call rate under 0.9 or an allele frequency lower than 5% in any tissue. A gene was considered expressed in a sample if its read count was greater than or equal to 6. Genes that were expressed in fewer than 10 of the samples in a tissue were removed for the eQTL analysis in that tissue. To correct for varying degrees of admixture in the African American subjects, we used the first three principal components of the genotyping data provided by the GTEx consortium and included these in our eQTL model. We used R R MatrixEQTL package [Shabalin, 2012] to calculate eQTLs with an additive linear model that included age, sex and ethnic background, as well as the first three genotype PCs, as covariates:

*Expression* ~ *Genotype* + *Age* + *Sex* + *Ethnic Background* + *PC*1_*genet*_ + *PC*2_*genet*_ + *PC*3_*genet*_ + *∊*

We tested for association between gene expression levels and SNPs both in *cis* and *trans*, where we defined *cis*-SNPs as those within 1MB of the transcription start site of the gene based on mapping using Bioconductor R biomaRt package [Durinck et al., 2009]. P-values were adjusted for multiple testing using Benjamini-Hochberg correction for *cis*- and *trans*-eQTLs separately and only those with adjusted p-values less than 0.2 were used in subsequent analyses.

### Community identification

For each tissue, we represented the significant eQTLs as edges of a bipartite network linking SNPs and gene nodes. To identify highly connected communities of SNPs and genes in the eQTL networks, we used the R condor package [Platig et al., 2016], which maximizes the bipartite modularity [Barber, 2007]. As recursive cluster identification and optimization can be computationally slow, we calculated an initial community structure assignment on the weighted, gene-space projection, using a fast unipartite modularity maximization algorithm [Blondel et al., 2008] available in the R igraph package [Csardi and Nepusz, 2006], then iteratively converged on a community structure corresponding to a maximum bipartite modularity.

The bipartite modularity is defined in Eq. (1), where *m* is the number of links in the network, *Ã_ij_*is the upper right block of the network adjacency matrix (a binary matrix where a 1 represents a connection between a SNP and a gene and 0 otherwise), *k*_*i*_ is the degree of SNP *i*, *d*_*j*_ is the degree of gene *j*, and *C*_*i*_, *C_j_* the community indices of SNP *i* and gene *j*, respectively.

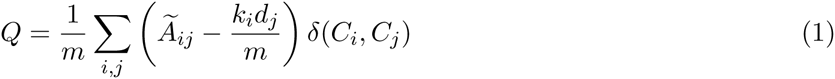

### SNP core score calculation

We defined a SNP’s core score as the SNP’s contribution to the modularity of its community. Specifically, For SNP *i* in community *h*, its core score, *Q_ih_*, is defined by Eq. (2). To normalize SNPs across communities, we accounted for community membership in our downstream testing (Eqns. 3 and 4), which better accounts for community variation compared to the normalization method used in [Platig et al., 2016].

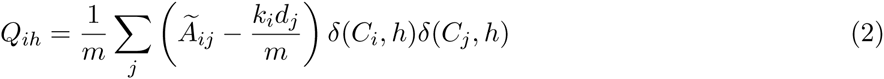

### Gene Ontology functional category enrichment

We extracted the list of genes within each community in each tissue-specific network, and then used the R GOstat package [Falcon and Gentleman, 2007] to perform a tissue-by-tissue analysis of the over-representation of Gene Ontology biological processes within each community. Our reference set consisted of all the genes present in the corresponding tissue-specific network. Communities were considered significantly enriched in a given category if the FDR-adjusted p-value was *<* 0.05.

### GWAS Analysis

We downloaded the NHGRI-EBI GWAS catalog (Accessed 08 DEC 2015, version v1.0) from the EBI website (https://www.ebi.ac.uk/gwas). We filtered out associations with *P*-values greater than 10^*−*^8. We then compared the distribution of SNP core scores between GWAS-associated SNPs from the NHGRI-EBI catalog and those not associated with traits or diseases for each tissue-specific network using a likelihood ratio test (LRT). In our setting, the LRT assess whether a linear model that includes GWAS status (Eq. (4)) fits the observed data better than a linear model that doesn’t include this variable (Eq. (3)). As the distribution of SNP core scores (*Q*_*ih*_) is not uniform across communities, we added community identity as a covariate in the linear regression. In Eqns. 3 and 4, *Q*_*ih*_ is the core score of SNP *i* in community *h*, *n* the number of communities in the tissue, *I*(*GW AS* = 1) an indicator function equal to 1 if the SNP is associated to a trait or disease in GWAS and equal to 0 otherwise, and *I*(*C*_*k*_ = 1) an indicator function equal to 1 if the SNP belongs to community *k* and equal to 0 otherwise.

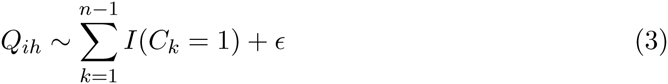

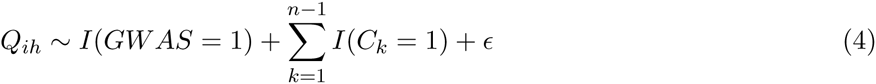

To control for linkage disequilibrium between SNPs, we generated lists of SNPs falling into the same LD block using the plink2 –blocks option and a 5MB maximum block size. In each community, for each LD block, we extracted the median of *Q*_*ih*_ for GWAS SNPs and non-GWAS SNPs, and used these values as input in the linear regressions.

### RegulomeDB category definition

We downloaded RegulomeDB data from the RegulomeDB website (http://www.regulomedb.org/downloads) and annotated all SNPs in the eQTL networks [Boyle et al., 2012]. We redefined RegulomeDB categories to exclude status as a previously reported eQTL, as all SNPs in our tests were eQTLs. Updated categories are described in Supplementary Table S4.

### Chromatin state category definition

We downloaded the genome-wide core 15 state model chromatin state data from the Roadmap Epigenome Project website (http://www.roadmapepigenomics.org/) for the eight tissues for which data are available: adipose subcutaneous, artery aorta, fibroblast cell line, esophagus mucosa, heart left ventricle, lung, skeletal muscle, and whole blood [Roadmap Epigenomics Consortium et al., 2015]. The fifteen chromatin states are active TSS (TssA), Flanking active TSS (TssAFlnk), Transcribed at a gene’s 5’ and 3’ end (TxFlnk), Strong transcription (Tx), Weak transcription (TxWk), Genic enhancers (EnhG), Enhancers (Enh), ZNF genes and repeated regions (ZNF/Rpts), Constitutive heterochromatin (Het), Bivalent/Poised TSS (Tss-Biv), Flanking bivalent TSS/enhancers (BivFlnk), Bivalent enhancers (EnhBiv), Repressed PolyComb (ReprPC), Weak repressed PolyComb (ReprPCWk), and Quiescent (Quies).

### RegulomeDB & Chromatin State Enrichment Analyses

For regulomeDB and chromatin state analyses, we calculated enrichment in each functional category among either global or local hubs using a logistic regression model [Kudaravalli et al., 2009] which allows for covariates:

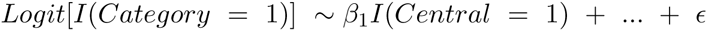

where *I*(*Category* = 1) is an indicator function equal to 1 if the SNP belongs to the functional category and equal to 0 otherwise, and *I*(*Central* = 1) is an indicator function equal to 1 if the SNP is central (in the top quartile of core scores or *>* 10 for degree), and equal to 0 otherwise. The odds ratios are estimated by *exp*(*β*_1_). We used all SNPs in the each tissue-specific network as background.

Similar to the calculation of enrichment in GWAS SNPs, we included indicators of community as a covariate when computing enrichment in regulomeDB and Roadmap Epigenomics Project categories among high core score SNPs. To control for the gene density around a SNP, which can impact the number of *cis*-associations, we added a covariate corresponding to the number of genes within 1Mb of a SNP.

Using the same method, we studied the enrichment in each chromatin state among trans-eQTLs for each tissue:

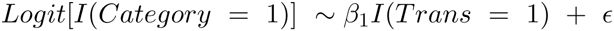

where *I*(*Trans* = 1) is an indicator function equal to 1 if the SNP is a *trans*-eQTL and equal to 0 otherwise. We used all SNPs with a MAF greater than or equal to 5% as background.

## Author Contributions

All authors conceived the study; MF, JNP, and JP analyzed the data; MF, JP, and JQ interpreted the results; MF, JP, and JQ drafted the initial manuscript. All authors contributed to the reviewing and editing of the manuscript. All authors read and approved the final manuscript.

## Acknowledgments

This work was supported by grants from the US National Institutes of Health, including grants from the National Heart, Lung, and Blood Institute (5P01HL105339, 5R01HL111759, 5P01HL114501, K25HL133599), the National Cancer Institute (5P50CA127003, 1R35CA197449, 1U01CA190234, 5P30CA006516), and the National Institute of Allergy and Infectious Disease (5R01AI099204). Additional funding was provided through a grant from the NVIDIA foundation. This work was conducted under dbGaP approved protocol #9112.

## Supplementary Figures

**Figure S1:**
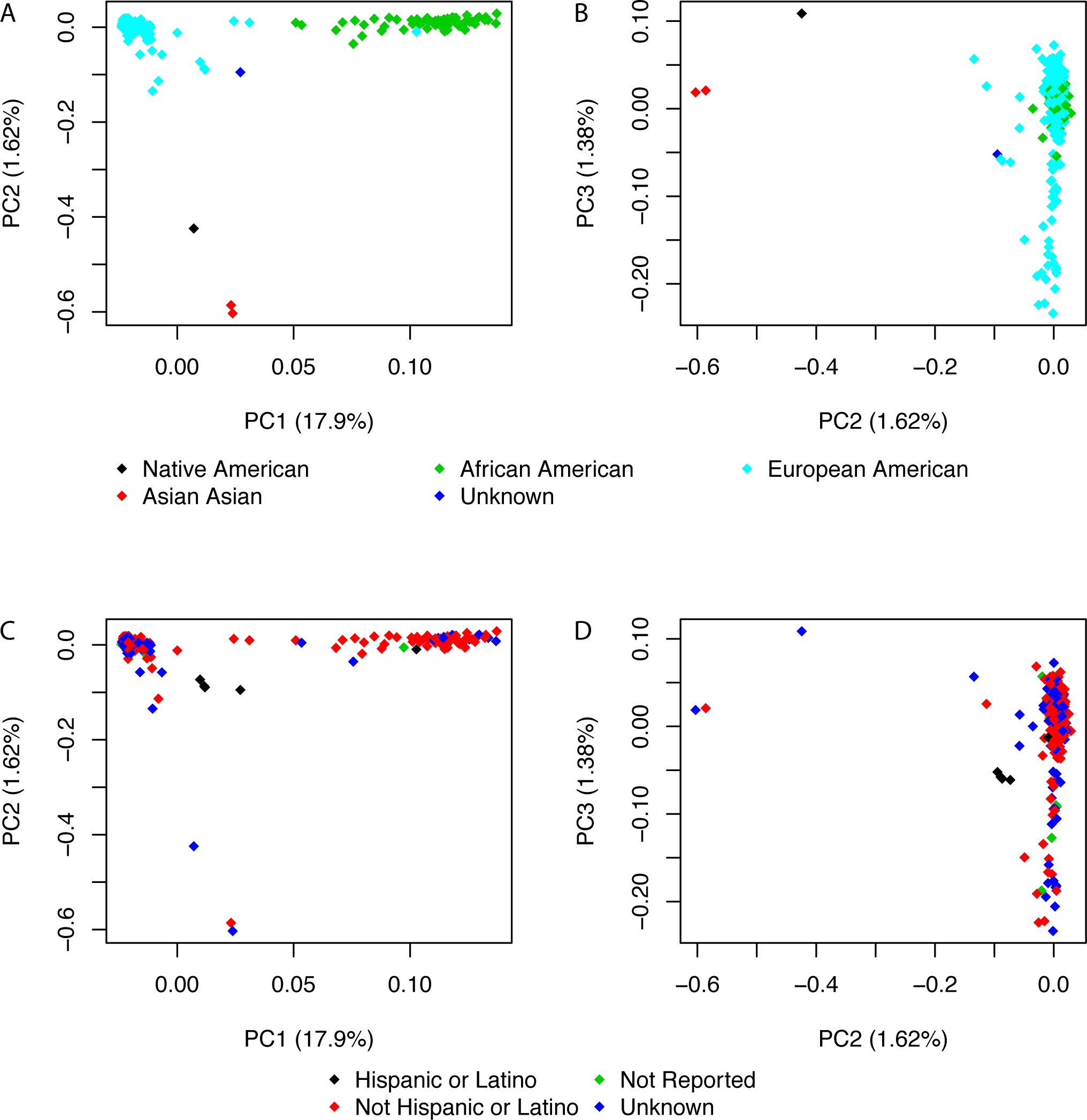
Related to Figure 1. Principal component analysis of genotyping data. **A.** and **C.** PC1 vs. PC2. **B.** and **D.** PC2 vs. PC3. **A.** and **B.** Colors represent population. **C.** and **D.** Colors represent ethnicity as reported by donor, family/next of kin, or medical record abstraction.

**Figure S2:**
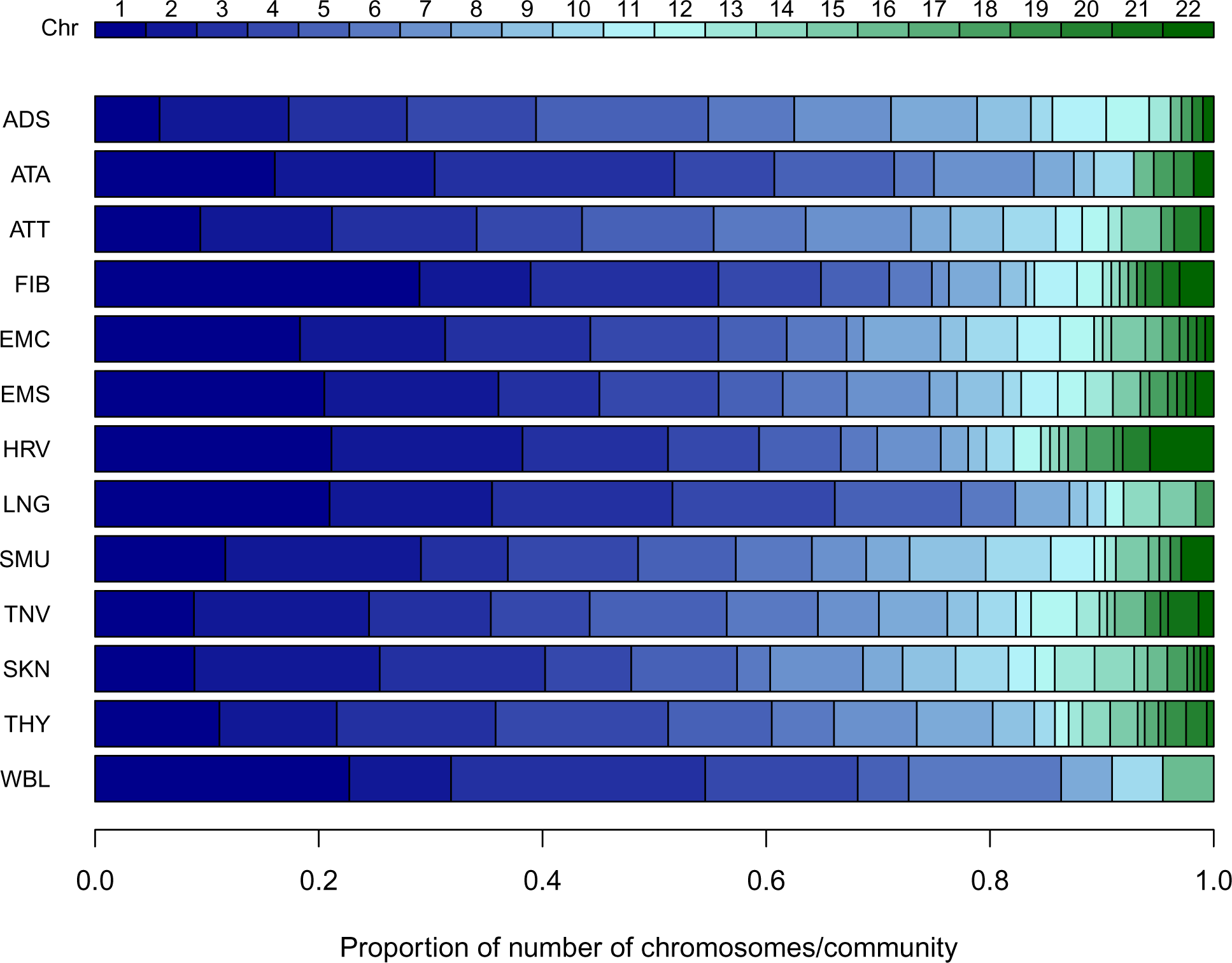
Related to Figure 3. Distribution of number of chromosomes by community for each tissue. Each barplot represent the proportion of communities that include SNPs and genes from each autosome.

**Figure S3:**
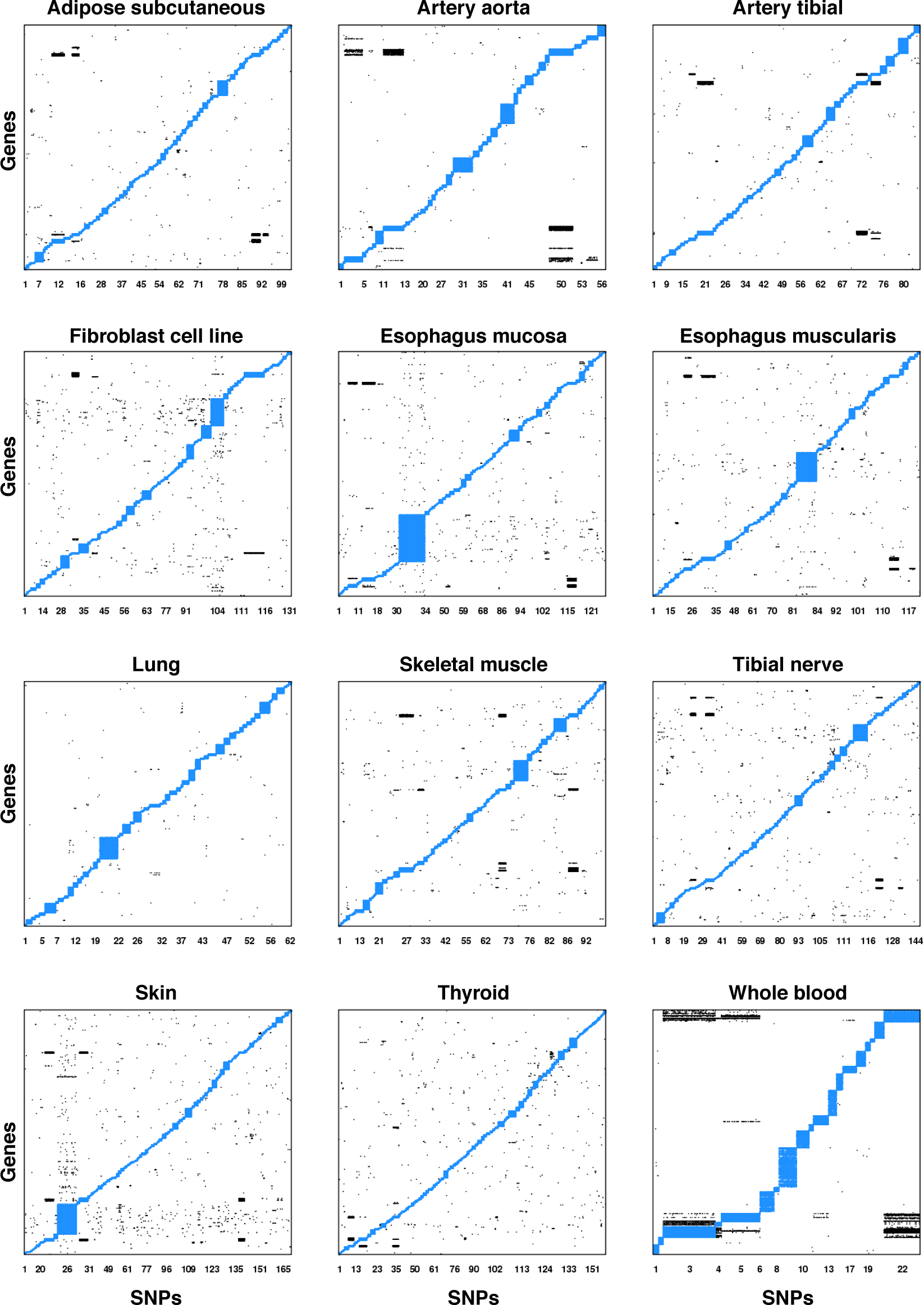
Related to Figure 3. Structure of communities within each eQTL network. Each edge is represented by a point. Intra-community edges are plotted in blue, inter-community edges in black.

**Figure S4:**
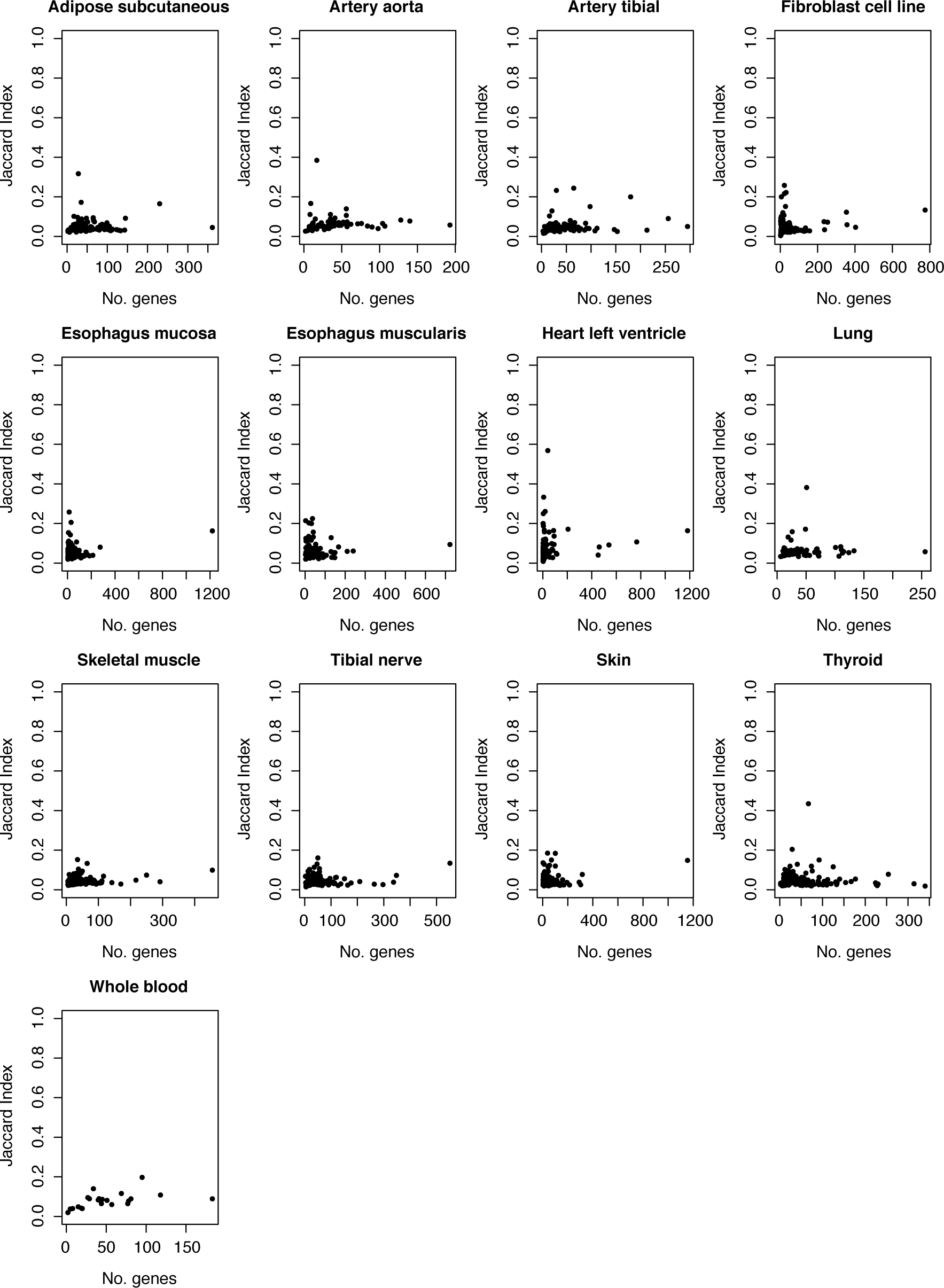
Related to Figure 2. Similarity of genes between eQTL network communities and gene expression correlation clusters. The correlation-based dendrogram was cut so the number of expression clusters and eQTL communities were equal. For each community, only the highest Jaccard Index value is given.

**Figure S5:**
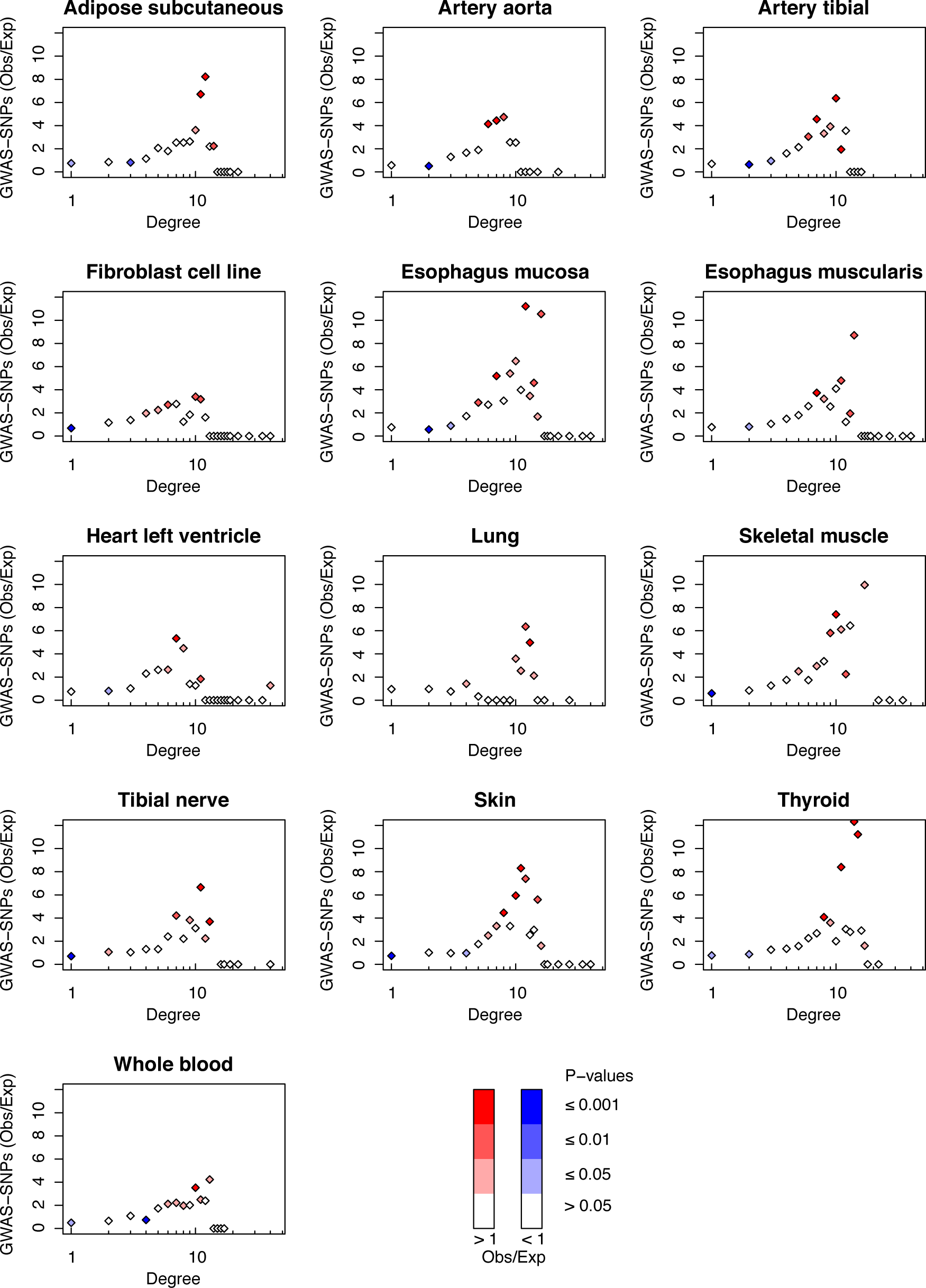
Related to Figure 6. Ratio of observed vs. expected number of SNPs associated to all traits or diseases by GWAS studies depending on their network degree for each tissue-specific network.

**Figure S6:**
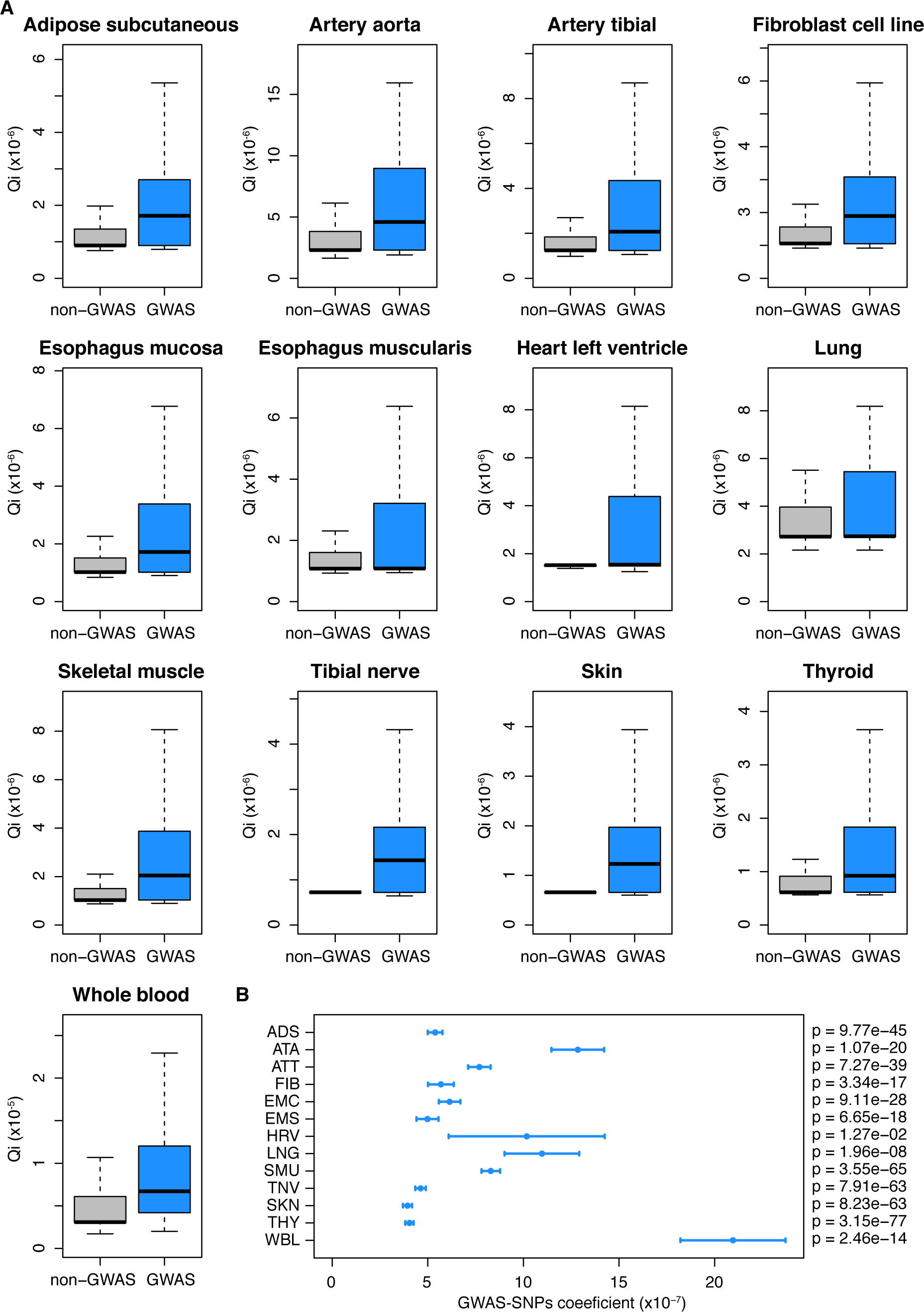
Related to Figure 6. GWAS-SNPs are enriched in high core score. A. Distribution of core-scores for SNPs associated (in blue) or not (in gray) to traits or diseases by GWAS studies for all tissues. B. Coe?cient for the GWAS term in the linear regression model. Bars indicate the standard error for the coe?cient. *P*-values indicated on the right were obtained using the likelihood ratio test described in the Methods.

## Supplementary Tables

**Table S1:** Related to Figure 1. Comparison of our results (at a FDR *<* 0.05) with GTEx *cis-eQTL* list.

| Tissues | GTEx | Our_analysis | Common eQTLs | Proportion common |
| --- | --- | --- | --- | --- |
| Adipose subcutaneous | 1,311,216 | 599,385 | 445,659 | 74.35% |
| Artery aorta | 878,091 | 446,775 | 313,000 | 70.06% |
| Artery tibial | 1,245,107 | 520,345 | 388,842 | 74.73% |
| Fibroblast cell line | 1,315,975 | 438,548 | 330,338 | 75.33% |
| Esophagus mucosa | 1,118,435 | 484,212 | 350,936 | 72.48% |
| Esophagus muscularis | 1,019,564 | 481,827 | 339,745 | 70.51% |
| Heart left ventricle | 620,472 | 285,283 | 201,318 | 70.57% |
| Lung | 1,095,502 | 483,266 | 355,454 | 73.55% |
| Skeletal muscle | 1,124,399 | 523,434 | 382,387 | 73.05% |
| Tibial nerve | 1,491,673 | 630,254 | 466,921 | 74.08% |
| Skin | 1,450,025 | 617,307 | 445,664 | 72.19% |
| Thyroid | 1,592,982 | 691,333 | 525,681 | 76.04% |
| Whole blood | 1,060,536 | 397,941 | 298,012 | 74.89% |

Table S2: Related to Figure 4. GO biological processes enrichment for each community in each tissue. cf. GeneOntology biological processes.csv file.

Table S3: Related to Figure 3. Association with traits and diseases of SNPs and genes from heart left ventricle community 86 and involved in cellular respiration biological processes. cf. list snps genes heart left ventricle 86.csv file.

**Table S4.** Related to Figure 5. Definition of functional classes and correspondence with regulomeDB categories.

| Class | RegulomeDB categories | Description |
| --- | --- | --- |
| A | 1a or 2a | TF binding + matched TF motif + matched DNase footprint + DNase peak |
| B | 1b or 2b | TF binding + any motif + DNase footprint + DNase peak |
| C | 1c or 2c | TF binding + matched TF motif + DNase peak |
| D | 1d or 3a | TF binding + any motif + DNase peak |
| E | 1e or 3b | TF binding + matched TF motif |
| F | 1f or 5 | TF binding + DNase peak |
| G | 4 | TF binding or DNase peak |
| H | 6 | Motif hit |
| I | 7 | No Information |

Table S5: Related to Figure 5. Enrichment in regulomeDB classes for central SNPs. cf. centrality regulomeDB enrichment.csv file.

Table S6: Related to Figure 5. Enrichment in epigenomic roadmap chromatin states for central SNPs. cf. centrality epigenomic roadmap enrichment.csv file.

Table S7: Related to Figure 5. Odds ratio and p-values measuring enrichment in each chromatin state for *trans*-eQTLs in each tissue. cf. transeQTLs epigenomic roadmap enrichment.csv file.

